# Endogenous viral elements constitute a complementary source of antigens for personalized cancer vaccines

**DOI:** 10.1101/2023.03.23.533908

**Authors:** Christian Garde, Michail A. Pavlidis, Pablo Garces, Emma J. Lange, Sri H. Ramarathinam, Mateo Sokač, Kirti Pandey, Pouya Faridi, Johanne Ahrenfeldt, Shanzou Chung, Stine Friis, Daniela Kleine-Kohlbrecher, Nicolai J. Birkbak, Jens V. Kringelum, Birgitte Rønø, Anthony W. Purcell, Thomas Trolle

**Affiliations:** Evaxion Biotech A/S, Dr Neergaards Vej 5F, Hørsholm, Denmark; Department of Biochemistry and Molecular Biology & Infection and Immunity Program, Biomedicine Discovery Institute, Monash University, Clayton, Victoria, Australia; Department of Molecular Medicine, Aarhus University Hospital, Aarhus, Denmark; Department of Clinical Medicine, Aarhus University, Aarhus, Denmark; Department of Medicine, School of Clinical Sciences, Monash University, Clayton, Victoria, Australia

**Keywords:** Endogenous retrovirus, personalized cancer vaccine, immunotherapy, neoantigen, T-cell epitopes, biomarker, immunopeptidomics

## Abstract

Personalized cancer vaccines (PCVs) largely leverage neoantigens arising from somatic mutations limiting their application to patients with relatively high tumor mutational burden (TMB). This underscores the need for alternative antigens to design PCVs for low TMB cancers. To this end, we substantiate endogenous retroviral elements (EVEs) as tumor antigens through large-scale genomic analyses of healthy tissues and solid cancers. These analyses revealed that the breadth of EVE expression in tumors stratify checkpoint inhibitor treated melanoma patients into groups with differential overall and progression-free survival. To enable the design of PCVs containing EVE-derived epitopes with therapeutic potential, we developed a computational pipeline, ObsERV. We show that EVE-derived peptides are presented as epitopes on tumors and can be predicted by ObsERV. Preclinical testing of ObsERV demonstrates induction of sustained poly-functional CD4+ and CD8+ T-cell responses as well as long-term tumor protection. As such, EVEs may facilitate and improve PCVs especially for low-TMB patients.

## INTRODUCTION

Neoantigen-based therapies designed to prime or amplify tumor-specific T-cell responses hold great promises for effective cancer immunotherapy^1,2^. With the regulatory approval of immune checkpoint inhibitors in multiple cancer types, immunotherapy has become standard of care for several advanced solid cancers^3^. Further development has demonstrated that augmented efficacy can be achieved through the combination of immunotherapies^4,5^. Next-generation sequencing of cancerous tumors has enabled the systematic discovery of patient-specific neoantigens and advanced immunotherapy to the stage of personalized medicine^6,7^. The design of personalized cancer vaccines (PCVs) has to date predominately focused on the identification of cancer-specific epitopes arising from somatic mutations. However, this limits the therapeutic scope of PCVs to cancers with high tumor mutational burden (TMB) as a considerable number of mutations are needed to select efficacious neoantigen targets. The hunt for novel efficacious tumor antigen sources was sparked to broaden and potentially enhance the application of PCVs^8–12^. Endogenous viral elements (EVEs), which include endogenous retroviruses (ERVs), account for remnants of ancient infectious viruses that have integrated their DNA into mammalian genomes. Under homeostatic conditions, expression of EVE genes in healthy tissues is largely considered silenced or kept at low levels through strict epigenetic regulation^13^. However, it has been documented that, in pathological conditions such as cancer and HIV-1 infection, loss of transcriptional regulation effectively leads to upregulated expression of EVE genes^14,15^. EVEs may thus constitute a source of tumor antigens to be harnessed by immunotherapy including PCVs^9^. Research has indicated that T-cell mediated immune recognition of human EVE (hEVE) peptides in healthy subjects is virtually absent, while HIV-1 and cancer patients mount functional T-cell responses against hEVE peptides^15–17^. A more recent study has used DNA barcode-based peptide-MHC (pMHC) multimers to confirm the absence of CD8+ T-cell-based recognition of hEVE peptides in several healthy subjects, and strikingly demonstrated hEVE peptide-specific CD8+ T-cell recognition in cancer patients^18^. To complement retrospective clinical findings, preclinical studies have demonstrated that vaccination with the EVE peptides, AH1 and p15E, confers tumor protection accompanied by EVE-specific T-cell responses^19–22^ and that EVE-targeting T cells may mediate anti-tumor effects induced by neoantigen immunotherapies^23^. These data show that EVE-specific T-cell responses are detectable in preclinical and clinical settings and highlight their potential for tumor control. Clinical intervention studies with TCR-transduced T-cells targeting ERVs are in the early stage of testing (NCT03354390), but there has to the authors’ knowledge not been conducted any clinical vaccine studies that leverage EVE-derived epitopes.

In the current study, EVEs are further substantiated as a source of tumor antigens. From large scale genomic analyses of healthy tissues and solid cancer biopsies we show that the number of expressed EVEs is cancer type specific with no correlation to the number of somatic variants (TMB), thereby suggesting that EVEs could augment the design of cancer vaccines. We developed a bioinformatics pipeline, ObsERV, for the design of PCVs comprising EVE-derived peptides. We confirm that EVE-derived peptides are presented on tumors as MHC ligands using state-of-the-art immunopeptidomics and that they can be predicted by ObsERV. Furthermore, ObsERV-designed therapies protected mice from tumor establishment and induced both specific CD4+ and CD8+ T-cell responses.

## RESULTS

### EVE expression in healthy tissue can inform vaccine target selection

Expression of a vaccine target in a healthy tissue may raise concerns about reduced efficacy due to central tolerance or safety risks from induction of autoreactive immunity^24^. While EVE genes at large have been considered silenced under normal conditions, there are reports of EVE expression in healthy tissue and their implications to host physiology. To define a set of EVEs, which can safely be used in vaccine development, we investigated their expression in human healthy tissues through RNA-seq analysis of GTEx deposited samples. Interestingly, some EVEs were sporadically expressed while others constituted tissue-specific signatures, as demonstrated by the 2D projection of the EVE expression profiles (Figure 1A). The tissues clustered well based on their EVE expression profile (Figure 1A and 1B), and some tissues had very distinct EVE expression profiles, e.g. muscle, testis, liver, and pancreas (Figure 1C). For each tissue except the immune privileged testis, the fraction of samples expressing a given EVE was computed (here referred to as the tissue fraction). If the highest tissue fraction exceeded 0.05, then the given EVE was considered expressed in healthy tissue otherwise it was labelled a potential cancer-specific target. To qualify the cut-off of the EVE expression levels, the EVE tissue fractions were compared to the well described melanoma antigen gene (MAGE) family. The MAGE family is considered restricted to immune privileged tissues and is upregulated in several solid cancers^25–29^. Members of the MAGE family have been tested as therapeutic vaccine targets in the clinic with acceptable safety profiles^25^. The MAGE family is thus a suitable comparator for cancer specificity. The highest tissue fractions of the cancer-specific EVE group were comparable or lower than those of the MAGE family, indicating that cancer-specific EVEs would be safe targets for vaccine development (Figure 1D).

**Figure 1.**
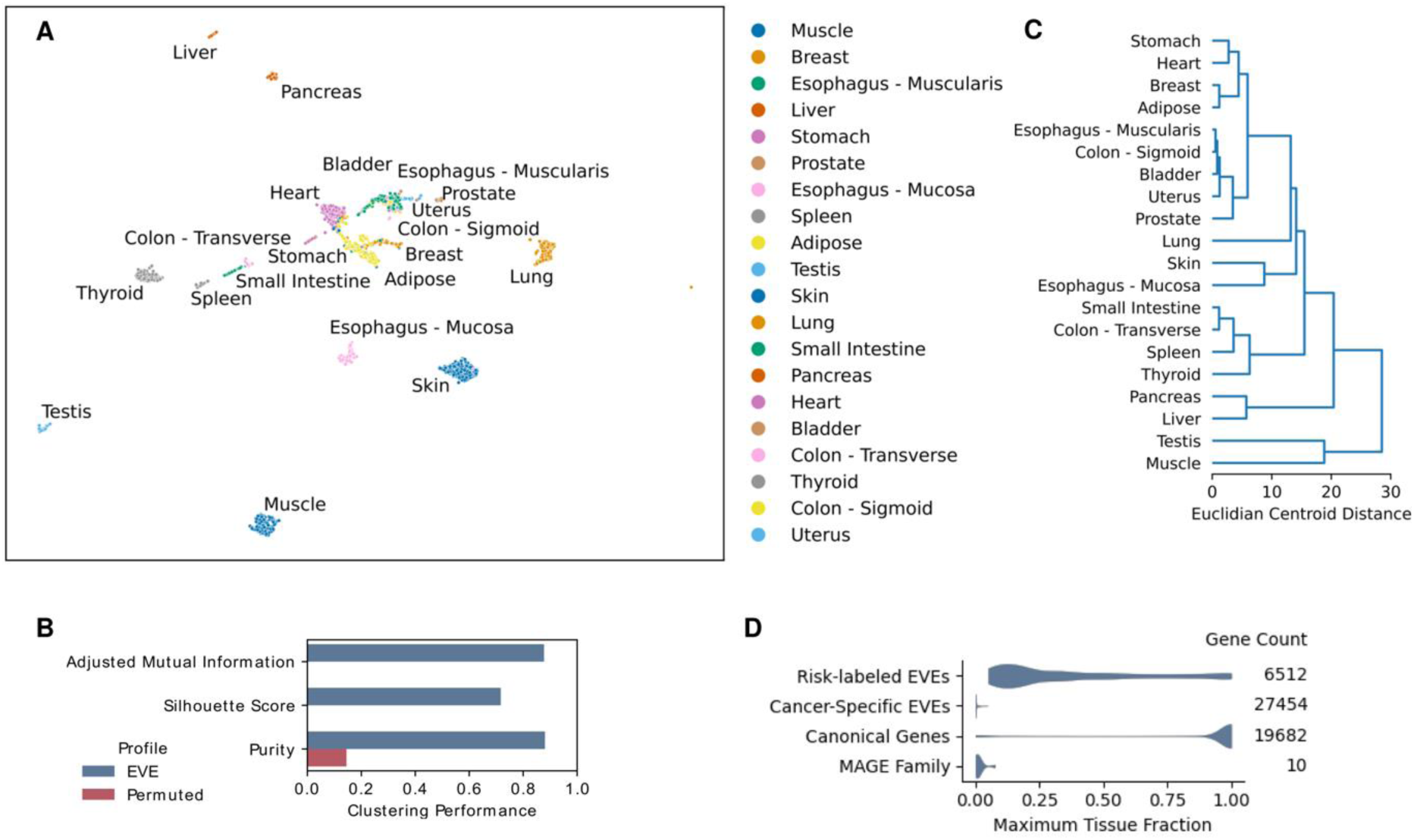
EVE expression across human tissues. (A) Uniform Manifold Approximation and Projection (UMAP) of EVE expression profiles in human tissues. (B) Clustering performance of the projected EVE expression profiles and a baseline established by permutation of tissue labels. (C) Agglomerative hierarchical clustering of UMAP tissue centroids summarizes the differences in tissue expression signatures. (D) Distribution of genes based on the fraction of tissue samples supporting their expression (TPM>1). The EVE genes are grouped based on expression in healthy tissues excl. immune privileged testis. Canonical protein-coding genes are shown as comparator along with a segregated group comprising the cancer-specific MAGE family.

### EVEs constitute a complementary antigen source for cancer vaccine development

Next, we investigated whether targeting EVEs could augment the design of cancer vaccines by serving as a complementary antigen source. Matched TCGA (The Cancer Genome Atlas) deposited RNA-seq and whole-exome sequencing (WES) of tumor biopsies from different solid cancer types were analyzed for EVE expression levels and somatic variants (Figure 2A, 2B, and 2C). We observed that the number of EVE genes expressed in the tumor (henceforth referred to as the EVE burden) differed across cancer types. Some cancers expressed relatively few cancer-specific EVEs, e.g. bladder cancer, while others displayed considerably higher EVE burdens, e.g. esophageal cancer and stomach adenocarcinoma. Notably, the EVE burden does not correlate with the mutational burden when comparing cancer types nor for patients belonging to the same cancer type (Figure 2D). To complement this, the fraction of patients with EVE burdens exceeding different thresholds was mapped out to further elucidate the feasibility of designing EVE-based PCVs in different cancer types (Figure 2E). To compare this with the coverage of a shared EVE vaccine product, we investigated the fraction of patients whose tumors express a shared set of cancer-specific EVE genes (Figure 2F). This provides an indication for the number of EVE antigens that could enter a shelf product if all its targets were required to be expressed by the tumor. The fraction of patients would naturally increase if the shelf product contained several targets and only a subset of them is required to be present in the tumor.

Overall, EVEs appear able to facilitate the design of PCVs for several cancer indications by augmenting the pool of cancer epitopes. In contrast, development of a shared vaccine product based on cancer-specific EVEs appears restricted to a more limited set of cancer types.

**Figure 2.**
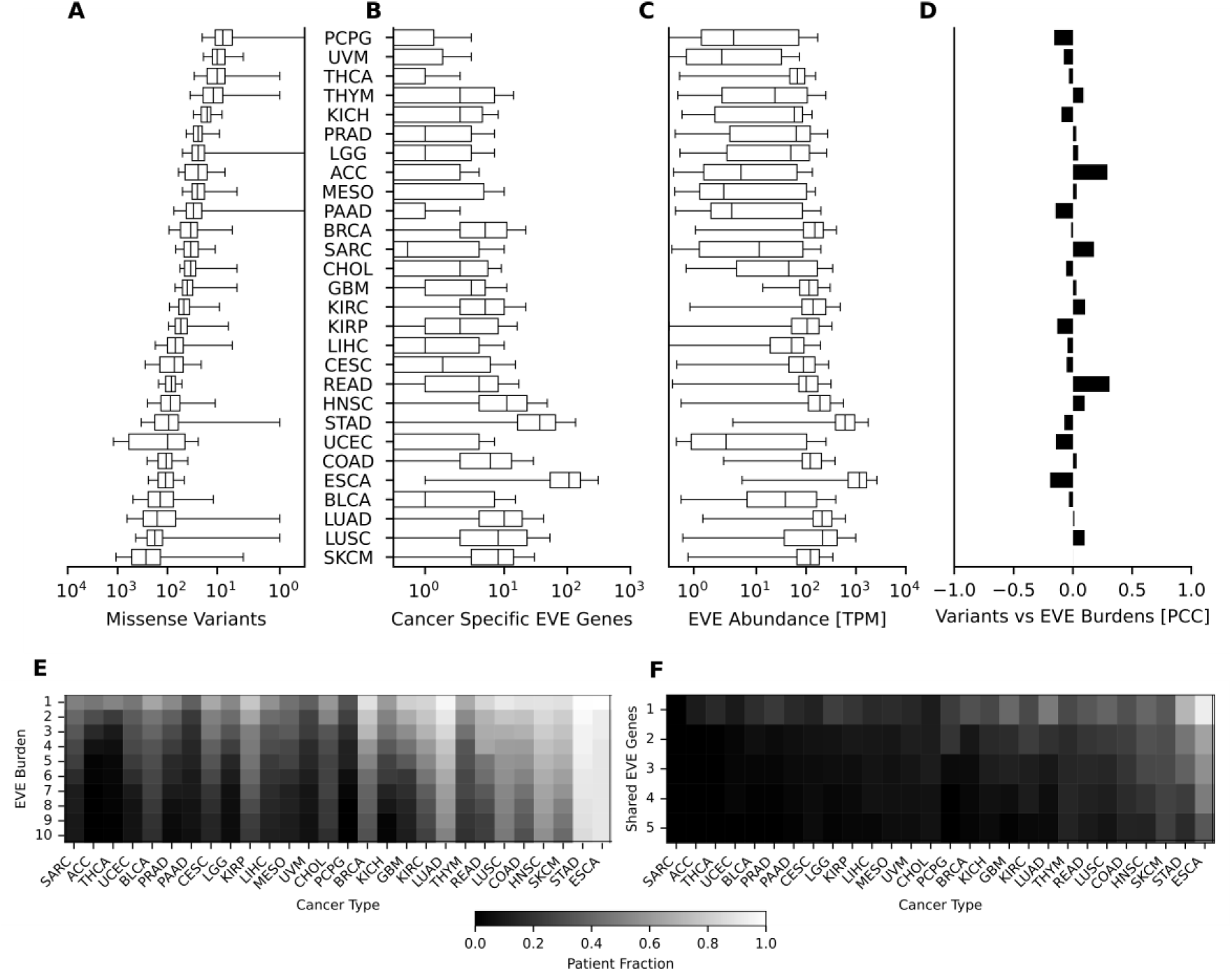
Pan-solid cancer analysis of EVE antigen burden and tumor mutational burden. (A) Distributions of the number of missense somatic variants. (B) Distributions of cancer-specific EVE burden. (C) Distributions of EVE transcript abundance in units of transcripts per million (TPM). (D) Correlation between number of missense variants and EVE burden. None of the Pearson correlation coefficients (PCC) were found statistically significant (all p-values > 0.05). Cancer types are abbreviated by the TCGA project and have the same ordering across panels A through D. TCGA project abbreviations definitions are provided in the Table S7 (E) Heatmap depicting the fraction of patients with EVE burden exceeding different thresholds to map the feasibility of PCVs. (F) Fraction of patients whose tumor express a shared combination of EVE genes to map the feasibility of a shelf vaccine product.

### The EVE burden stratifies cancer patients receiving checkpoint inhibitor therapy

EVEs have been linked to cancer development and immune suppression^30,31^. To supplement these findings, we explored the correlation between EVE expression and clinical outcome of cancer patients. Specifically, the investigation focused on whether the EVE burden could stratify cancer patients into groups with differential overall and progression-free survival. Three studies of metastatic melanoma patients receiving checkpoint inhibitor therapy were included, namely Liu et al.^32^, van Allen et al.^33^ and Riaz et al.^34^. The dataset was filtered to include only patients with pre-therapy tumor biopsies characterized by RNA-seq and WES and a healthy tissue biopsy characterized by WES resulting in 107 patients for the survival analyses. The mutational burden was represented by the number of missense somatic mutations supported with a variant allele frequency of at least 5% based on analysis of the matched tumor/healthy WES, and the RNA-seq was analyzed for EVE expression levels to compute the EVE burden. To ensure balanced downstream analyses, study-wise medians were used to assign group labels to the patients based on their mutational burden and EVE burden (TMB-High/Low and EVE-High/Low). The studies were combined to a melanoma ensemble to augment the number of patients. Interestingly, the EVE-Low group had longer overall survival compared to the EVE-High group (Figure 3A) and showed trends of longer progression-free survival (Figure 3D). The EVE based stratification did not display differential survival for the patients assigned to the low mutation burden group (Figure 3C and 3F). In contrast, patients assigned to the high mutational burden group showed enhanced differential overall and progression-free survival between the EVE burden groups (Figure 3B and 3E). To complement this, we also analyzed 245 bladder cancer patients treated with anti-PDL1 therapy with tumor samples characterized by RNA-seq and WES^35^. We did not find any differences in overall survival between groups defined by the median EVE burden overall nor in the TMB-High group (Figure S2).

**Figure 3.**
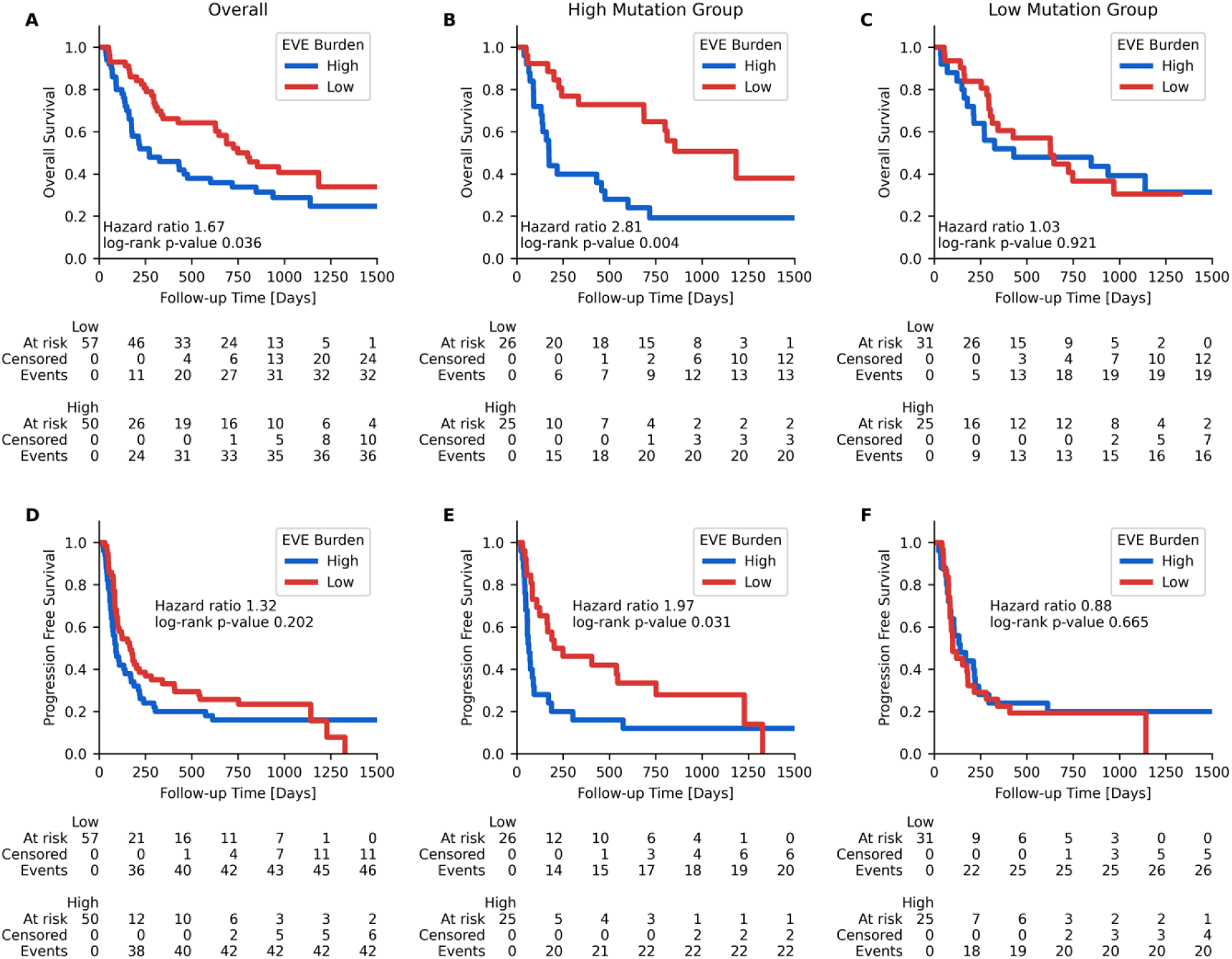
Survival analysis of checkpoint inhibitor treated melanoma patients by the EVE burden. (A-C) Overall survival analysis of melanoma patients using the EVE burden: (A) all patients, (B) high mutation subgroup patients, (C) low mutation subgroup of patients. (D-F) Progression-free survival analysis of melanoma patients using the EVE burden: (D) all patients, (E) high mutational subgroup patients, (F) low mutation subgroup of patients.

### EVEs are presented as MHC ligands on tumors and constitute a potential source of antigens for personalized cancer vaccines

To investigate the feasibility of using EVE-derived tumor antigens in PCVs, we developed an *in silico* bioinformatics pipeline, ObsERV. ObsERV relies on RNA-seq data and MHC ligand predictions to identify and select EVE peptide sequences that are likely to be presented on MHC molecules (Figure 4A). The first step in the ObsERV pipeline is to leverage RNA-seq data to quantify EVE expression levels in the tumor sample. For all expressed EVE genes, MHC class I (MHCI) ligand predictions are generated for all 8-11mer peptides and MHC class II (MHCII) ligand predictions are generated for all 15mer peptides. MHC ligand predictions are combined, and MHC ligand hotspots are identified (see Methods). Finally, highest scoring ligand hotspots from each EVE are ranked based on the integrated MHC presentation scores, generating a list of candidate EVE-derived peptide sequences which can be selected for vaccine production and administration.

To verify that EVEs are translated, processed, and presented as MHC ligands, we performed immunopeptidomics on CT26 and B16F10 mouse cancer cells and tumor samples. For CT26, three immunopeptidomics datasets were generated using unstimulated cells, IFNy-stimulated cells and *in vivo* grown tumors. A total of 43679 unique peptide sequences without post translational modifications (PTMs) were identified across these three datasets (Figure S3A). For B16F10, two immunopeptidomics datasets were generated using IFNy-stimulated cells and *in vivo* grown tumors. Unstimulated B16F10 cells were not analyzed as previous studies have reported poor MHC presentation on unstimulated B16F10 cells^36^. A total of 7078 unique sequences without PTMs were identified across the two datasets (Figure S3G). The datasets were further filtered, keeping only peptides with a length of 8-11 amino acids, a PEAKS confidence value above 20 and expression levels above 1 TPM. Following these filtering steps, a total of 23826 MHCI ligands remained in the CT26 dataset, of which 89 originated from EVE sequences (Figure S3B). For the B16F10 dataset, a total of 4749 MHCI ligands passed the filtering criteria with 19 originating from EVEs (Figure S3H). We next clustered the MHC ligands by their sequence motifs and generated sequence logos for each cluster. For CT26, the sequence logos matched known motifs for H2-Dd, H2-Kd and H2-Ld (Figure S3D, S3E and S3F), and for B16F10, the sequence logos could be matched to H2-Db and H2-Kb, as expected (Figure S3J and S3K).

Finally, we set out to evaluate the predictive performance of the tools employed by ObsERV to predict surface presented MHCI ligands derived from expressed EVEs and canonical proteins, defined as transcripts with TPM above 1. Three methods for selecting EVE ligands were benchmarked; 1) EVE expression levels alone, 2) sequence-based MHC ligand presentation predictions alone and 3) a combination of 1 and 2. For comparison, a naïve approach where MHC ligands were selected at random was also included. To ensure an unbiased assessment of the predictive performance of the ObsERV tools, the CT26- and B16F10 datasets described above were further filtered to remove sequences that were included in the training dataset of the MHC ligand prediction tool utilized by ObsERV. The CT26 benchmark dataset consisted of 54 EVE-derived ligands and 10673 endogenous ligands, while the B16F10 benchmark dataset included 18 EVE-derived ligands and 1645 endogenous ligands. For each benchmark dataset, negative data points were added by randomly sampling peptide sequences from other EVEs/proteins with TPM>1 at a 30:1 negative to positive ratio. As expected, the integrated model of expression and ligand prediction outperforms expression and ligand predictions alone as well as the random model. For B16F10, the performance on EVE ligands was overall higher than on the canonical protein derived ligands, although this could be due to the low number of EVE ligands included in this dataset. For the CT26 datasets, the expression levels alone also performed better on the EVE ligand subset compared to the canonical protein ligand subset, which was not seen with the other methods, where the performance values were similar across ligand types.

Overall, this analysis shows that EVEs are expressed in mouse cancer models and that EVE-derived peptides are processed and presented by MHC molecules on the cell surface. We demonstrated that the EVE-derived ligands can be predicted using *in silico* methods already employed for prediction of canonical protein derived MHC ligands, indicating that EVE-derived peptides are a viable source of tumor antigens for PCVs.

**Figure 4.**
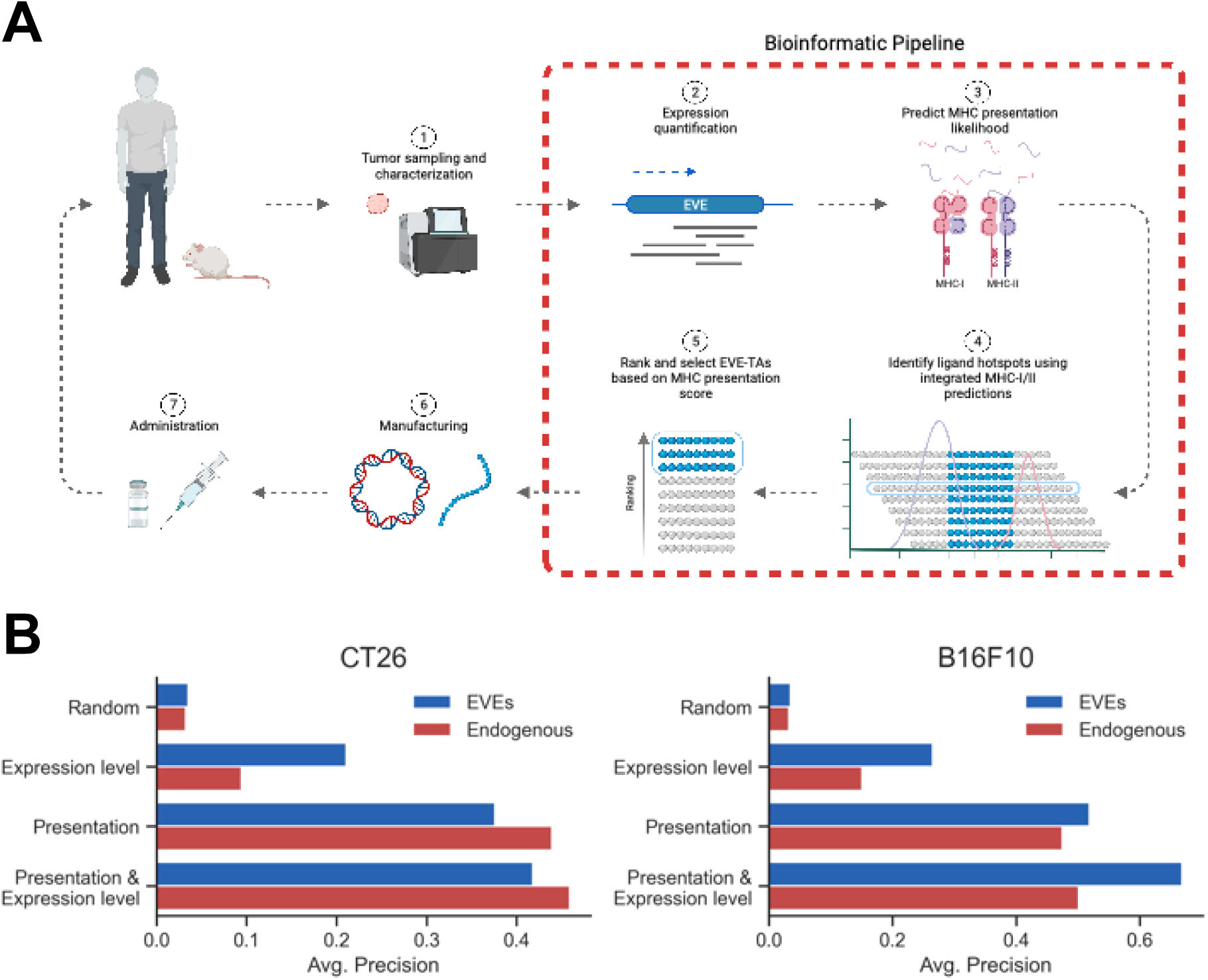
Pipeline overview and evaluation of predictive performance. (A) Flow chart for designing EVE-based immunotherapies. A tumor sample is collected for characterization using RNA-seq. For all expressed EVEs, the likelihood of MHC ligand presentation is predicted for all 8-11mer peptides, and a ligand hotspot is selected based on an integrated MHCI and MHCII presentation score. The EVE hotspots are then ranked based on the combined presentation scores and EVE expression levels, resulting in a final prioritized list of EVE-derived peptide candidates. The top scoring EVE-derived peptides are selected for manufacturing and administration. (B) Benchmark of features for selecting EVE-derived peptides in the CT26 and B16F10 mouse cancer models. Bars show the average precision score of the four features evaluated in the benchmark. Blue bars show the predictive performance on EVE-derived MHC ligands and red bars show performance on MHC ligands from endogenous proteins.

### *In silico* designed EVE-based cancer vaccines induces strong anti-tumor effect and robust T-cell responses in preclinical cancer models

We set out to investigate whether EVEs are relevant therapeutic cancer targets for personalized cancer vaccines using preclinical models of colon carcinoma (CT26) and melanoma (B16F10). First, the EVE expression levels in *in vivo* grown CT26 and B16F10 tumors were quantified using RNA-seq. Next, using ObsERV, we designed vaccines specific to each tumor based on the EVE expression levels and predicted MHC ligands. For CT26, the *in silico* design comprised the 13 top ranked EVE peptides encoded into a pTVG4 plasmid DNA (pDNA) (from hereon: ObsERV_CT26). Of note, the well described H2-Ld-restricted EVE epitope SPSYVYHQF (AH1) derived from the gp70 Envelope (Env) protein of the Murine Leukemia Virus (MuLV) was selected by ObsERV (O3_CT26). For B16F10, the vaccine design included the top 5 ranked EVE epitopes formulated as a peptide vaccine (from hereon: ObsERV_B16F10). The H2-Kb-restricted minimal epitope KSPWFTTL derived from p15E, the transmembrane subunit of the Env protein of MuLV, was selected by the ObsERV platform as an ObsERV_B16F10 epitope (O2_B16F10). The amino acid sequences of the vaccine peptides and their computational features are shown in Tables S3 and S4. We started our investigations in the CT26 model. Groups of mice were vaccinated with ObsERV_CT26, mock pDNA or a pDNA encoding predicted neoantigens (NeoAgs_CT26) previously shown to induce anti-tumor effect^37^ in one-week intervals starting two weeks prior to subcutaneous (s.c.) inoculation of CT26 cancer cells (Figure 5A). Notably, the ObsERV_CT26 vaccine prevented tumor establishment in almost all vaccinated mice, whereas treatment with mock pDNA did not prevent the development of tumors (Figures 5B and 5C). These results highlight that the ObsERV-designed vaccine confer strong tumor protection in this model. We then assessed the immune responses elicited by the ObsERV_CT26 vaccine using the IFNγ ELISPOT assay. Re-stimulation of isolated splenocytes with the ObsERV_CT26 pool yielded robust IFNγ responses, while re-stimulation with each vaccine peptide individually identified O3_CT26, O6_CT26 and O8_CT26 as the main drivers of immunogenicity (Figure 5D). Re-stimulation with the minimal AH1 peptide encoded in the highly immunogenic vaccine epitope O3_CT26 revealed that AH1 underlies the strong immune recognition of O3_CT26 (Figure 5D). Immune analysis in splenocytes from mock DNA-immunized mice revealed strong IFNγ responses against the ObsERV_CT26 pool, with peptide O3_CT26 being the sole contributor (Figure 5D). In contrast, naïve mice showed no immunity against the ObsERV_CT26 pool (Figure 5D), indicating that immune recognition of the pool and specifically AH1 in the mock group was related to tumor presence. Overall, these data show that the ObsERV-designed vaccine induced strong anti-tumor immune responses in the CT26 model.

To gain a deeper understanding of the immune responses induced by ObsERV_CT26 vaccination, we characterized the functional phenotype of the vaccine-specific T cells using the Intracellular Cytokine Staining (ICS) assay. The ObsERV_CT26 vaccine elicited polyfunctional CD8+ and CD4+ T cells reactive to the vaccine pool (Figure 5E and F). Furthermore, re-stimulation with each of the ELISPOT-identified immunogenic peptides revealed that peptides O3_CT26 and O6_CT26 were strong inducers of IFNγ and TNFα production within splenic CD8+ T cells, in line with reports on the ability of AH1 to drive CD8+ T-cell responses (Figure 5E)^21^. Vaccine epitopes O6_CT26 and O8_CT26 induced polyfunctional CD4+ T-cell reactivity (Figure 5F). Finally, immune recognition of the ObsERV_CT26 pool in the mock group was driven by polyfunctional CD8+ T cells recognizing the peptide O3_CT26 (Figure 5E). Taken together, our results underscore that the ObsERV-designed vaccine led to elicitation of a balanced, multifunctional CD4+ and CD8+ T cell response in the CT26 model.

Motivated by the strong immunogenicity elicited by the ObsERV_CT26 vaccine, we examined whether these responses were long-lasting. To that end, a subset of ObsERV_CT26-vaccinated, tumor-free mice along with age-matched, treatment-naive mice were interrogated for T-cell immune recognition of AH1 epitope 79 days after the CT26 tumor challenge, corresponding to 63 days after the last immunization (Figure 5G). Using the MHC class I dextramer methodology, we observed persistent AH1-directed CD8+ T-cell recognition in the blood of the vaccinated mice, whereas naïve mice did not harbor a specific response (Figure 5H). To investigate whether ObsERV_CT26 immunotherapy conferred durable tumor protection, vaccinated mice were re-challenged with CT26 cells 85 days after the primary tumor challenge. In contrast to the untreated mice that developed tumors, mice that had received ObsERV_CT26 vaccine demonstrated a long-lasting anti-tumor effect, as almost all of them remained tumor-free following the CT26 tumor re-challenge (Figures 5I and 5J).

Further immune analyses demonstrated that mice vaccinated with ObsERV_CT26 preserved 80% of the immune reactivity to the ObsERV_CT26 pool measured in the primary study and sustained their capacity to recall strong IFNγ responses against the same vaccine epitopes identified as immunogenic previously (Figure 5K).

Collectively, the data demonstrate that an *in silico* designed, EVE-targeting vaccine leads to long-lasting anti-tumor effect and persistent EVE-specific T-cell immunity in the CT26 cancer model.

**Figure 5.**
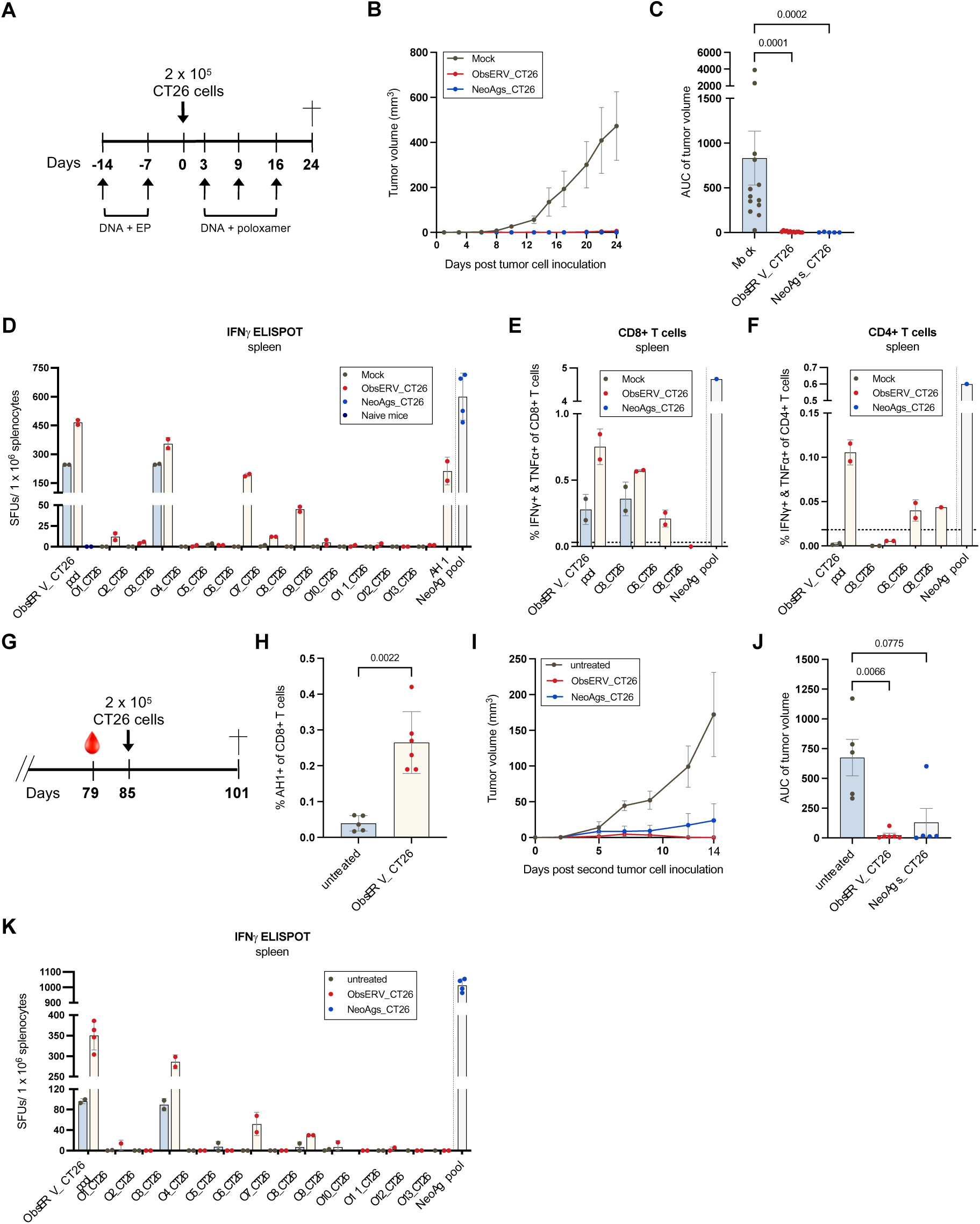
*In silico* designed EVE-based vaccine induces long-lasting tumor protection and T-cell immunity in the CT26 colon carcinoma model. (A) Study outline of the *in vivo* tumor study. (B) Group mean tumor growth curves (in mm^3^) ± standard error of the mean (SEM). (C) Area under the tumor growth curve (AUC) for individual mice by group. Mean ± SEM. (D) Pooled bulk splenocytes (n = 6-7 mice) were tested on IFNγ ELISPOT assay for immune reactivity to the ObsERV_CT26 pool and the individual vaccine peptides (run in technical duplicates). Mean +/-Standard Deviation (SD). Stars indicate the immunogenic vaccine peptides carried on to the re-stimulation and ICS. (E) and (F) Pooled bulk splenocytes (n = 6-7 mice per group) were re-stimulated with the ObsERV_CT26 pool and the individual vaccine peptides identified as immunogenic in IFNγ ELISPOT in (D) followed by ICS for detection of polyfunctional CD8+ (E) and CD4+ T cells (F) (run in duplicates). Mean +/-SD. Dotted line represents background levels of cytokine production in unstimulated samples. (G) Study outline of the CT26 tumor re-challenge study. (H) Percent presence of AH1-specific CD8+ T cells in tail vein blood of ObsERV_CT26-vaccinated mice and age-matched untreated mice 79 days after the primary CT26 tumor challenge. Blood was stained with MHC class I dextramers comprising H2-Ld loaded with the restricted epitope AH1 epitope (SPSYVYHQF). Mean +/-SD. (I) Group mean tumor growth curves (in mm^3^) ± SEM. (J) AUC for individual mice by group ± SEM. (K) Pooled bulk splenocytes (n = 4-6 mice per group) were tested on IFNγ ELISPOT assay for immune reactivity to the ObsERV_CT26 pool and the individual vaccine epitopes (run in duplicates). Mean +/-SD. Experiment was run twice. *Statistics*: Matt-Whitney t-test (H), Kruskal-Wallis test with Dunn’s multiple comparisons correction (C, J).

We then turned to investigate whether EVE-based vaccines confer tumor protection in the B16F10 melanoma model. Groups of mice were vaccinated with ObsERV_B16F10 comprising the top five ObsERV-selected EVE epitopes as peptides adjuvanted with polyinosinic:polycytidylic acid (polyI:C) or with polyI:C alone. As a positive control, mice were vaccinated with a combination of a minimal epitope derived from the Tyrosinase-related protein 2 (Trp2) and three publicly described B16F10-specific neoantigens shown to drive strong anti-tumor effect (TAA_NeoAgs_B16F10)^22,37^. Vaccination commenced two weeks prior to s.c. B16F10 tumor challenge and was repeated in one-week intervals (Figure 6).

Vaccination with ObsERV_B16F10 led to strong B16F10 tumor growth delay, as the majority of mice developed significantly smaller tumors on day 18 compared to the polyI:C alone group (Figure 6B and C). Notably, the ObsERV_B16F10-driven anti-tumor effect was comparable to that attained by the TAA_NeoAgs_B16F10 treatment. Immune analysis in isolated splenocytes from ObsERV_B16F10-vaccinated mice revealed robust T-cell responses against the vaccine epitope pool (Figure 6D). Combined, these data highlight the tumor-protective effect of ObsERV_B16F10 PCV, mirrored in the elicitation of vaccine-specific T cell responses.

We then asked whether the EVE-based PCV induced changes in the immune composition of the Tumor Microenvironment (TME). Staining of single cell suspensions of isolated tumor digests from the ObsERV_B16F10 group revealed considerable enrichment of CD45+ immune cells compared to tumors from mice treated with polyI:C alone (Figure 6E). ObsERV_B16F10-treated tumors demonstrated a strong tendency for increased presence of intratumoral CD4+ T cells, in contrast to FoxP3+ CD4+ T-cell numbers that remained unchanged, thus suggesting that the enriched CD4+ T cells bore a non-regulatory phenotype (Figure 6F and 6G). Notably, ObsERV_B16F10 treatment led to strong infiltration of CD8+ T cells (Figure 6H), whereas no differences were noted in tumor-infiltrating NK cells (Figure 6I). Taken together, these experiments indicate that the ObsERV_B16F10-induced anti-tumor effect is reflected on a robust reshaping of the TME, with increased infiltration of T cells.

Collectively, our preclinical experiments suggest that EVEs can serve as relevant cancer antigens and be targeted in the context of *in silico* designed PCV, leading to strong and efficacious CD8+ and CD4+ T-cell responses.

**Figure 6:**
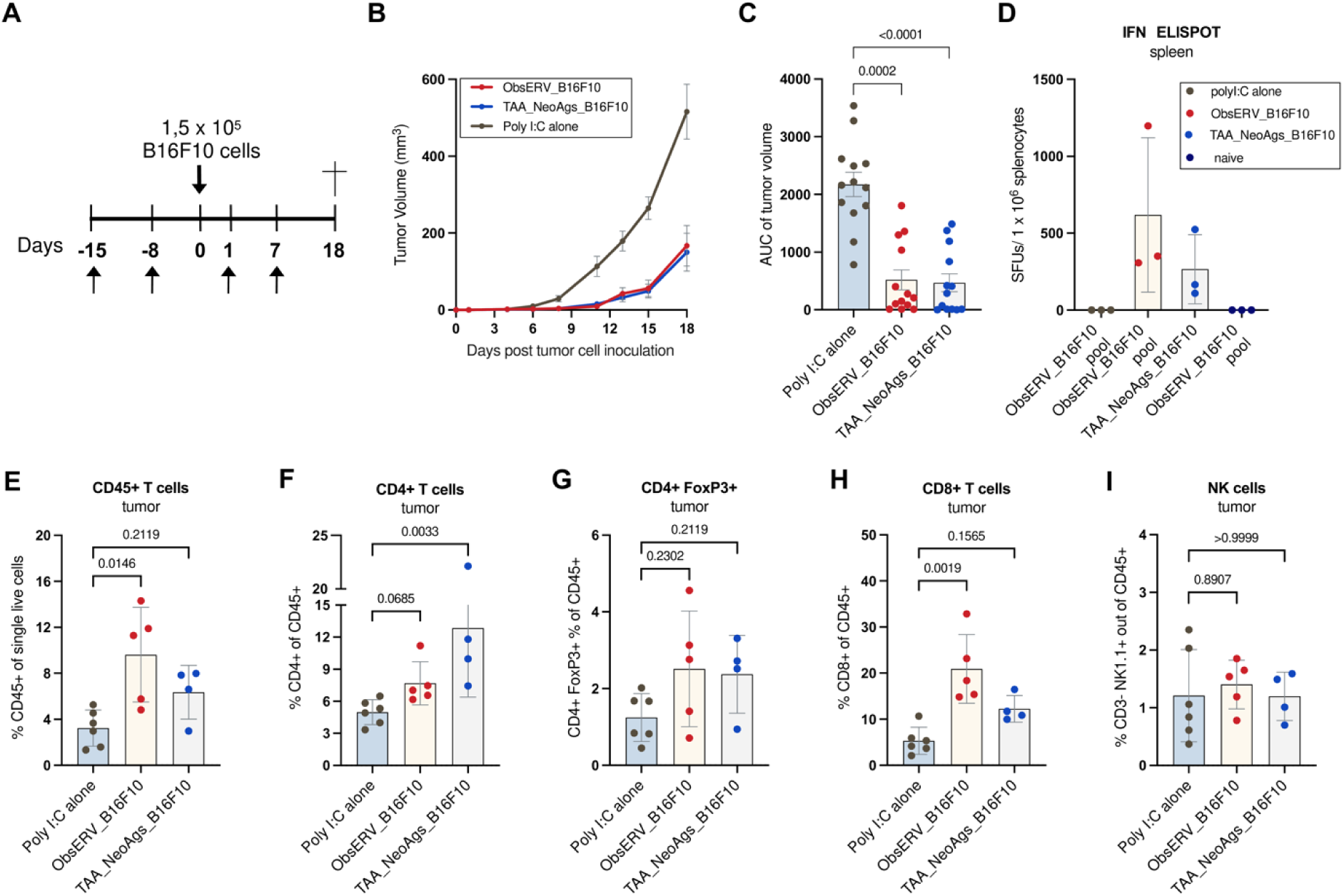
In silico designed EVE-based immunotherapy induces robust B16F10 tumor growth delay and intratumoral T-cell enrichment. (A) Study outline of the in vivo tumor study. (B) Group mean tumor growth curves in mm3 +/-SEM. (C) AUC for individual mice by group. Mean +/-SEM. (D) Splenocytes from individual mice (n=3 mice per group) were tested on IFNy ELISPOT assay for immune reactivity against the ObsERV CT26 pool (run in technical duplicates). Mean +/-SD. (E)-(I) Single cell suspensions of B16F10 tumor digests were analyzed for the presence of (E) immune cells (CD45+), (F) CD4+ T cells out of CD45+, (G) CD4+ FoxP3+ out of CD45+, (H) CD8+ T cells out of CD45+ and (I) NK cells (CD3-NK1.1+) out of CD45+ cells. Mean +/-SD. Statistics: Kruskal-Wallis test with Dunn’s multiple comparisons correction (C) (E-I)

## DISCUSSION

Additional tumor antigens are needed to expand the application of PCVs to more cancer patients especially those with low TMB. We set out to further substantiate EVEs as a source of tumor antigens to be leveraged by PCVs and developed a computational pipeline, ObsERV, for rational PCV design. First, we investigated healthy human tissues and found that EVE expression signatures clustered the samples by tissue, which is in line with recent reports^26,27^. Accumulating evidence suggests that some EVEs exert functions relevant for the host’s physiology^28,29^. The tissue-specific EVE expression may support this dogma. The analysis also revealed which EVEs are silenced in human tissue and may be considered safe targets for cancer vaccine development. Notably, the majority of EVEs were considered silenced in healthy tissue, while most of the canonical protein coding genes were expressed in at least one tissue. Next, a pan solid cancer analysis revealed that the EVE burden is cancer type specific and that it does not correlate with TMB. As such, EVEs may be used as a supplementary antigen source to pave the way for PCV designs for some low-TMB patients and select cancer types that are considered unsuitable target indications for neoantigen-based PCVs. Furthermore, a brief analysis suggests that a shelf vaccine product based of cancer-specific EVEs may be tractable for a limited set of solid cancers. To advance this, more research is warranted on selecting the optimal set of vaccine sequences to match a segment of cancer patients. EVEs are subject to epigenetic control and the pool of EVE antigens could potentially be further enhanced by co-administration of epigenetic modulators^39–41^. In extension, EVEs could be an interesting vaccine target for treating e.g. hematological malignancies such as acute myeloid leukemia where such epigenetic modulators already are an established line of therapy^42^.

Based on published melanoma cancer cohorts receiving checkpoint inhibitor therapy, survival analyses further support the relevance of EVEs in cancer. We find that the breadth of EVE expression specifically stratifies TMB-High labeled melanoma patients into groups with differential overall and progression-free survival. This stratification was not observed in a large bladder cancer cohort receiving checkpoint inhibitors, which might be explained by the fact that low numbers of EVEs were found to be expressed in both the TCGA bladder cancer cohort, as well as in the cohort of bladder cancer patients treated with anti-PDL1. The EVE burden could potentially be used in combination with TMB to improve the identification of cancer patients who would benefit from checkpoint inhibitor therapy. Smith et al. (2018)^43^ used overall survival analysis and identified an ERV signature in renal cell carcinoma that was linked to clinical outcome, supporting that EVEs can serve as clinical biomarkers. Furthermore, Ng et al. (2023)^44^ found that lung patients with ERV-specific antibodies displayed improved response to checkpoint inhibitor therapy, indicating that eliminating retroviral expressing cancer cells is beneficial for patient outcome. This aligns with our analyses on the melanoma patients, where the low EVE breadth group displayed longer survival as compared to the high EVE breadth group. Interestingly, *in vitro* studies of cell lines have associated the expression of ERVs to tumorigenesis^45^ and interactions with various oncogenes including *MYC* have been suggested^31,46^. *MYC* plays a key role in regulating cell growth and proliferation, and it is frequently found to be amplified in multiple different cancers, including melanoma^47,48^. Retroviral envelope proteins have also been reported to suppress the immune system and associate with the immunosuppressor TGF-ß^31^. TGF-ß has been associated with poor outcome of checkpoint inhibitor therapy^49^. Some EVEs may thus serve as a fallback mechanism of immune evasion and facilitate resistance to checkpoint inhibitor therapy.

Collectively, these genomic analyses of healthy and cancerous tissue support EVEs as therapeutic targets for PCVs. This motivated us to develop an *in silico* platform, ObsERV, for the design of PCVs based on EVE-derived peptides. We demonstrated that EVEs are presented by MHC class I on the cell surfaces of murine tumors, validating that EVE derived peptides are eligible targets for T-cell mediated therapy. These observations are in line with previous studies that have found that peptides from various transposable elements are presented on MHC molecules and may elicit immune responses^9,10,18,50^. ObsERV was found capable of predicting the EVE-derived MHCI ligands for both the CT26 and B16F10 murine models, motivating the advancement of ObsERV to preclinical testing.

Preclinical testing demonstrated that vaccination with ObsERV-selected EVE epitopes confers strong anti-tumor effect in two tumor models accompanied by robust, vaccine-specific T-cell responses. While cases of preclinical anti-tumor protection of endogenous retroviral therapy have been previously reported for select model antigens^19,20,21^, this is to our knowledge the first time that an *in silico* designed immunotherapy comprising EVE-derived epitopes led to tumor protection and strong, poly-functional T-cell responses. Encouragingly, the immunodominant CD8+ T-cell epitope AH1 from the gp70 Env protein of MuLV was selected by ObsERV as a top vaccine epitope. AH1 has been shown to drive strong CD8+ T-cell responses and to confer protection in the CT26 model^19–21,23^, but optimal efficacy has been demonstrate to rely on assisted presentation, either through co-delivery with CD4+ T-cell helper epitopes during priming or peptide pulsing of *in vitro* pre-activated Dendritic Cells (DCs)^19,51^. In our study, ObsERV_CT26 vaccination elicited strong AH1 immunogenicity, and AH1-specific CD8+ T cells were strongly functional and detectable months after the last vaccination. We hypothesize that the broad immunogenicity and the strong persistence of vaccine-specific T-cell responses months after last antigen exposure, which translated into rejection of tumor establishment upon secondary tumor challenge, are supported by the induction of vaccine-specific, Th1-polarized CD4+ T cells. Indeed, it is well documented that CD4+ T cells play a crucial role in potentiating anti-tumor CD8+ T-cell responses in preclinical models^52^ by enabling their expansion and differentiation during the priming phase^53^ and by regulating memory CD8+ T-cell differentiation^54^. This highlights the importance of incorporating CD4+ T-cell epitopes in the vaccine design to achieve a robust and durable T-cell response profile and validates the ObsERV ranking strategy that seeks to elicit both CD4+ and CD8+ T-cell responses.

ObsERV_B16F10 vaccination led to significant tumor growth delay and vaccine-specific immune responses of comparable magnitude to the TAA_NeoAgs_B16F10 pool encompassing well characterized antigens. Interestingly, one of the ObsERV_B16F10 epitopes, O2_B16F10, derived from p15E, the transmembrane subunit of the Env protein of MuLV, and encompasses a described CD8+ T-cell minimal epitope. Similarly to AH1, p15E-based vaccination confers protection against B16F10 tumors or other C57Bl/6-syngeneic tumors^55^ only when of CD4+ T cells are concomitantly engaged, for example, through agonistic anti-CD40/CD40L antibodies^51,56^. The lack of knowledge on the individual ObsERV_B16F10 epitope immunogenicity and the reactive T-cell subsets that underlie the observed vaccine pool recognition limits our ability to conclude on the induction of p15E-specific T cells and the nature of the immune response. However, the tumor characterization experiment indicated significant TME immune remodeling in ObsERV_B16F10-treated mice, indicating a potential involvement of CD8+ T cells and, to a lesser extent, non-regulatory CD4+ T cells. This model reaffirms the ability of the ObsERV platform to select immunogenic EVE epitopes that confer strong tumor protection.

Collectively, we have demonstrated that EVEs can be potent anti-tumor targets that can be leveraged to facilitate and potentially enhance PCV development.

## METHODS

### EXPERIMENTAL MODELS AND SUBJECT DETAILS

#### Mouse cell lines

The BALB/c syngeneic colon cancer cell line CT26 (#CRL2638) was purchased from ATCC and cultured in R10 medium prepared from RPMI (Gibco #72400-021) supplemented with 10% heat inactivated fetal calf serum (FCS, Gibco # 10500-064) at 37 °C and 5% CO2 as per supplier’s instructions. The murine melanoma cell line B16F10 derived from a C57BL/6 mouse (ATCC, cat# CRL-6475) was purchased from ATCC and grown in DMEM (Sigma, cat# D6546) supplemented with 10% Fetal Bovine Serum (Gibco, cat# 10500-064) and 1% Glutamax (Gibco, cat# 35050-061). Cells were grown to 60-70% confluency, trypsinized and washed 2x in serum free RPMI in preparation for inoculation in mice.

#### Animal studies

Animals were maintained in the University of Copenhagen, Panum 10.3, Animal Facility, Copenhagen, Denmark or at the animal facility at Evaxion Biotech, Hørsholm, Denmark. All experiments were conducted under license 2017-15-0201-01209 from the Danish Animal Experimentation Inspectorate in accordance with the Danish Animal Experimentation Act (BEK nr. 12 of 7/01/2016), which is compliant with the European directive (2010/63/EU).

For RNA-seq and immunopeptidomics analysis, 6-8 week old female BALB/cJRj and C57BL/6JRj SPF mice were acquired from Janvier Labs (France). The mice were acclimated for one week before initiation of experiments. BALB/cJRj and C57BL/6JRj mice were subcutaneously (s.c.) inoculated with 2 x 10^5^ CT26 or B16F10 tumor cells, respectively, on the left flank in a volume of 100 μl serum free RPMI. Upon establishment, tumors were measured three times a week using a digital caliper and tumor volumes, *V*, were calculated using the following formula:

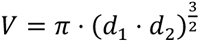

where *d*_1_ and *d*_2_ are the orthogonal diameters of the tumor. B16F10 tumors were harvested on day 15 and CT26 tumors were harvested on day 19-21. Tumors were immersed in RNAlater for RNA-seq or snap frozen in an ethanol bath on dry ice and maintained at -80 °C before immunopeptidomics analysis.

For tumor challenge studies, mice from the different experimental groups were distributed across different cages to avoid potential cage effects. For the CT26 study, mice were vaccinated weekly in left and right tibialis anterior muscles (i.m.) with 100 μg of research-grade DNA for a total of five immunizations. Vaccination commenced two weeks prior to s.c. CT26 cell inoculation (defined as study day 0). In the first two immunizations, 2 x 50 μL vaccine comprising DNA formulated in PBS was administered using Electroporation (EP) (BTX Agile pulse, VWR). In the last three immunizations, DNA was formulated with block co-polymer poloxamer 188 (gifted by BASF, Germany) to a final concentration of 3% in PBS and administered in 2 x 75 μL vaccine solution. Poloxamer has been shown to increase the longevity of the pDNA after injection and thereby increase antigen expression and exposure^58^. At the day of CT26 cell inoculation, 2 x 10^5^ of *in vitro* expanded CT26 cells were inoculated s.c. in the right flank of the mice. For the CT26 tumor re-challenge experiment, 2 x 10^5^ *in vitro* expanded CT26 cells were inoculated in the opposite (left) flank. Upon establishment, tumors were measured using the above-described formula. Mice were euthanized through cervical dislocation when the majority of tumors in the control groups reached the maximum allowed size of 15 mm diameter in either direction or upon reaching humane endpoints. For the B16F10 study, mice were vaccinated intraperitoneally (i.p.) with groups of *in vivo* grade peptides admixed with polyI:C [PolyI:C (HMW) VacciGrade™, Invivogen #vac-pic]. *In vivo* grade peptides comprising the ObsERV_B16F10 and TAA_NeoAgs_B16F10 groups were purchased from BioSynth (Netherlands), had purity ≥ 88% and were administered at 150 µg of each peptide per dose, except for peptide 01_B16F10 that was administered at 60 µg per dose. PolyI:C and was prepared according to manufacturer’s instructions and administered at 100 µg per mouse per dose. Vaccination commenced two weeks prior to s.c. B16F10 cell inoculation (defined as study day 0) and repeated weekly for a total of four immunizations. B16F10 cell inoculation (1,5 x 10^5^ cells per mouse), subsequent tumor measurement and mouse euthanization was conducted as in the CT26 study described above.

#### Generation of single cell suspensions of splenocytes

Suspensions of splenocytes were generated as described before^37^. Upon euthanization, spleens from mice with tumors representative of their group’s average tumor size were collected in cold RPMI supplemented with 10% FCS (from here on: R10), followed by processing to single cell suspensions via GentleMACS processing (Miltenyi Biotec, C-tubes #130-096-334 and Dissociater #130093-235) and passage through a 70 μm filter (Corning, CLS431751). Splenocytes were cryopreserved in FCS supplemented with 10% DMSO (Merck, #D8418).

#### Generation of B16F10 tumor digests

Tumor digests were generated as described before^37^. In brief, isolated tumors were dissociated into single-cell suspensions with a cocktail of tumor dissociation enzymes (Miltenyi Biotech #130-096-730) and filtered through 70 μm cell strainers according to the instructions of the manufacturer.

### METHOD DETAILS

#### RNA sequencing of mouse tumors

Total RNA was extracted from three CT26 tumors and three B16 tumors using the RNeasy Plus Mini Kit (Qiagen #74134) and shipped to GenomeScan (Leiden, The Netherlands) for sequencing. RNA-seq libraries were prepared using the NEBNext Ultra II Directional RNA Library Prep Kit for Illumina (NEB #E7760S/L) following the kit protocol. Briefly, mRNA was isolated from total RNA using oligo-dT magnetic beads, fragmented and used for cDNA synthesis. Sequencing adapters were ligated to the resulting cDNA and PCR amplified to create the final sequencing libraries. The libraries were sequenced on an Illumina NovaSeq 6000, generating ∼40M clusters of 150bp PE reads for each tumor sample.

#### Quantification of EVE expression

Reference genomes and associated GTF annotation files were downloaded from Ensembl^59^ for mouse (GRCm38.102) and human (GRCh38.104). Mouse- and human EVE nucleotide and amino acid sequences and GTF files were retrieved from the gEVE database^60^ v1.1. EVEs mapping to alt contigs not present in the Ensembl reference genomes were removed from the gEVE GTFs.

For both mouse and human, a combined reference + EVE GTF was created by concatenating the reference GTF files with modified gEVE GTF files where each feature field had been modified to “transcript” and “exon”. The combined GTF and reference genome fasta files were used to create STAR and RSEM indexes.

Raw fasta files for each tumor sample were preprocessed using cutadapt^61^ v1.18 and trimmomatic^62^ v0.38 to remove adapter sequences and poor quality bases. Preprocessed RNA reads were mapped to the GRCm38/GRCh38 reference using STAR^63^ v2.7.9a and transcript expression was quantified using RSEM^64^ v1.3.1. For biological replicates, combined expression profiles were calculated using the median TPM for each transcript. To ensure proper quantification, only samples with at least 1e7 reads assigned to protein-coding genes or EVEs were used in downstream analyses.

#### Immunoprecipitation of pMHC complexes

Cells were lysed in a lysis buffer containing 1% IGEPAL (Sigma), 50 mM Tris pH 8, 150 mM NaCl and protease inhibitors (cOmplete™, EDTA-free Protease Inhibitor Cocktail tablets, Roche Molecular Biochemicals) as described previously^65,66^. Lysates were cleared by centrifugation and MHC-peptide complexes were isolated using antibodies specific to the MHC molecule(s) of interest. Kd-, Ld- and Dd-associated peptides were selectively immunopurified for CT26 cell lines using the SF1.1.10, 28-14-8s and 34-1-2s antibodies, and a pan-MHCI approach was employed for the CT26 tumor and both B16 experiments where all three antibodies were mixed in equal proportions. The HLA-peptide complexes were then eluted using 10% acetic acid and subsequently subjected to molecular weight cut-off filters (small scale) or fractionated using a reversed-phase C18 end capped HPLC column (Chromolith SpeedRod, Merck), running a mobile phase buffer A of 0.1% trifluoroacetic acid (TFA) and buffer B of 80% acetonitrile (ACN) and 0.1% TFA. The peptides were separated using a gradient of 2-40% buffer B for 4 minutes and 40-45% for another 4 min. Fractions collected were then concentrated using centrifugal evaporation and reconstituted in 2% ACN in 0.1% formic acid (FA) prior to analysis by LC-MS/MS.

#### Data acquisition by LC-MS/MS

Reconstituted peptides were analyzed using a timsTOF Pro mass spectrometer, where reconstituted peptides were loaded onto an IonOpticks Aurora 25 cm C18 column using a nanoElute liquid chromatography system (Bruker Daltonics). The peptides were separated using a gradient of buffer B (0.1% FA in ACN) against buffer A (0.1% FA, 2% ACN in water) initially to 17% over 60 min then to 25% over 30 min then to 37% over 10 min with a flow rate of 300 nl/min. Data dependent acquisition (DDA) was performed with the following settings: mz range: 100-1700mz, capillary voltage: 1600V, target intensity of 30000, TIMS ramp of 0.60 to 1.60 Vs/cm2 for 166 ms.

#### Identification of MHC ligand sequences

LC-MS/MS data was searched with PEAKs Xpro v10.6 (Bioinformatic solutions) against the Uniprot annotated proteome^67^ (2020_06 release for mouse and 2021_04 release for human) appended with the gEVE database^60^ for the corresponding organism. The search was carried out using the following settings: Enzyme: none, parent mass tolerance 20 ppm, fragment mass tolerance 0.02 Da, Variable modifications: Deamidation (NQ), Oxidation (M). For the B16F10 datasets, peptides with C-terminal lysine- and arginine residues were filtered to remove potential tryptic peptide contamination.

#### Benchmarking of MHC ligand predictions

For each MHC ligand identified by immunopeptidomics, source protein/EVE expression levels were determined as described previously. MHC ligand predictions were generated for all relevant MHC alleles (H2-Dd, H2-Kd, H2-Ld for CT26 and H2-Db and H2-Kb for B16) using a proprietary MHC ligand prediction tool developed by Evaxion Biotech. As input, the tool was provided with the ligand sequences and the source protein/EVE expression values. As output, the tool calculated two MHC ligand probability scores based on: i) the amino acid sequence alone and ii) the amino acid sequence and the expression level.

To generate positive datapoints for the benchmark datasets, the MHC ligand sequences determined immunopeptidomics were filtered to only keep ligands that passed the following criteria: i) -log10(p) > 20, ii) peptide length 8-11, iii) derived from a source transcript/EVE with TPM>1 and iv) not included in the training dataset of the MHC ligand prediction tool.

Negative datapoints were generated by sampling peptide sequences from the Ensembl reference proteome and the gEVE database. Predictions were generated for all negative datapoints as described above. Negative datapoints were subject to the same filters as those applied to the positive datapoints. The final benchmark datasets were compiled by sampling negative datapoints at a 30:1 negative to positive ratio. For each datapoint in the benchmark datasets a random value was sampled from a uniform distribution between 0-1 using Numpy v1.23.5. Average precision scores were calculated using scikit-learn^68^ v1.1.3.

#### Selection of EVE-derived epitopes

EVE epitope sequences were selected by first predicting MHC ligands. Predictions were generated for all peptide sequences in the gEVE database of lengths 8-11 for MHCI molecules and of length 15 for MHCII molecules. For each EVE amino acid sequence, all unique 27mers were extracted and the best MHCI and -II ligands were determined based on predicted ligand probabilities. A combined MHC ligand score, *S*, was calculated for each 27mer using the following equation:

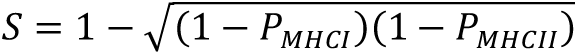

Where *P_MHCI_* and *P_MHCII_* are the ligand probabilities for MHC class I and -II respectively. The list of 27mers was sorted by their combined scores and filtered to keep only the highest scoring 27mer from each EVE and the top ranking 27mers from the resulting sorted- and filtered list were selected.

For preclinical testing with the CT26 cancer model, MHC ligands were predicted for the BALB/c MHC type, i.e., H2-Dd, H2-Kd, H2-Ld and H2-IAd as described above. The top 13 ranking EVE 27mer sequences were selected. For preclinical testing with the B16F10 cancer model, MHC ligands were predicted for the C57BL 6 MHC type, i.e., H2-Db, H2-Kb and H2-IAb as described above. The top 5 ranking EVE 27mer sequences were selected based on the MHC class I scores.

#### Generation of DNA plasmids encoding CT26 EVE epitopes

DNA sequences were generated for each EVE epitope using reverse translation through weighted random sampling of codons based on observed codon frequencies in the mouse genome. The EVE epitope DNA sequences were then combined with linker sequences encoding 10 amino acid glycine-serine linkers to form a DNA insert sequence. This process was repeated to generate 10 000 DNA insert sequences. The list of DNA sequences was then filtered to retain sequences with the lowest GC content and highest codon adaptation index (CAI). The GC content and CAI thresholds were defined so that at least 20% of the sequences were retained when each threshold was applied individually, and at least 10% of the sequences when applied together. The resulting list of DNA sequences was further filtered to remove any containing Kozak sequences. The predicted minimum free energy (MFE) of the corresponding RNA molecule to each remaining DNA sequence was calculated using ViennaRNA^69^ v2.4.18 and the best scoring insert was selected. A Kozak sequence and restriction sites were added to final DNA insert before it was manufactured and cloned into a pTVG4 DNA plasmid. The pTVG4 DNA vector was generated from the standard plasmid pUMVC3 acquired from Aldevron (North Dakota, USA, #4010) by cloning two copies of a 36-bp CpG-rich immunostimulatory DNA sequence (ISS) containing the 5’-GTCGTT-3’ motif downstream of the multi-cloning site. The EVE epitope containing pTVG4 DNA plasmid was upscaled by Aldevron. An empty pTVG4 DNA vector with no cloned EVE epitope DNA insert was also ordered to serve as a mock control.

#### Detection of immunogenic epitopes with IFNγ ELISPOT assay

T-cell reactivity against vaccine-encoded epitopes was measured *ex vivo* using the Enzyme-Linked ImmunoSPOT (ELISPOT) assay. In brief, 5 x 10^5^ of viable splenocytes suspended in R10 medium [RPMI 1640 (Fisher Scientific #72400021) + 10% FBS (Fisher Scientific #10500064)] were plated on an ELISPOT plate (Merck Millipore #MAIP4510) coated overnight with 5 μg/mL anti-murine IFNγ capture antibody (BD #51-2525KZ). Cells were stimulated with 5 μg/mL of synthetic vaccine-corresponding 27mer peptides, peptide pools or left unstimulated (R10 medium only). ObsERV_CT26-and ObsERV_B16F10-corresponding peptides used for *ex vivo* stimulations were purchased from BioSynth (Netherlands). Cells were stimulated for 20 hours at 37 °C and 5% CO_2_. IFNγ produced upon re-activation of peptide-specific T cells and captured on anti-IFNγ capture antibody was visualized via sequential addition of 2 μg/mL biotinylated anti-IFNy detection antibody (BD #51-1818KA), Streptavidin-Horseradish Peroxidase (HRP) (BD #557630) and AEC chromogen substrate (BD #551951) as per manufacturer’s protocol. After overnight drying, the ELISPOT plates were imaged in an ELISPOT reader (Cellular Technology, Ltd) and bound IFNγ was calculated as Spot Forming Units (SFUs) per 1 x 10^6^ splenocytes.

#### MHC class I dextramer staining for AH1-specific CD8+ T cells

To monitor T-cell immune recognition of vaccine epitopes in blood of mice, the MHC class I dextramer staining methodology was used. MHCI dextramers comprised H2-Ld molecules loaded with the restricted minimal peptide AH1 (SPSYVYHQF) derived from the MuLV gp70 protein and encoded by the vaccine epitope O3_CT26 (TYHSPSYVYHQFERRAKYKREPVSLTL). MHC class I dextramers were acquired from Immudex (#JG3294) and were fluorescently labelled with PE. In brief, 50 μL of tail vein blood was collected in EDTA-coated tubes (Sarstedt #20.1278.100) and then transferred in a 96 deep-well plate (Sigma #575653). Blood cells were then incubated with the FcR-blocking anti-murine CD16/32 antibody (Biolegend #101301) for 10 minutes at 4 °C. Prior to use, the dextramer reagent was centrifuged at 3300 g for 5 minutes. 10 μL of the dextramer reagent was added in each blood sample and incubated at 37 °C for 15 minutes. Cells were subsequently stained with fluorescently labelled antibodies specific for murine CD3e (FITC, Biolegend #100305), CD4 (PE-Cy7, BD #552775), and CD8 (BV786, BD #563332) at 4 °C for 30 minutes. Finally, cells were subjected to one-step fixation/red blood cell lysis (eBioscience #00-5333-57) before acquisition on FACS Celesta (BD). Determination of AH1-recognizing cells out of CD8+ T cells were determined using FlowJo Software (version 10.8.0). The gating strategy is presented in Figure S4A.

#### Stimulation and Intracellular cytokine staining (ICS) of epitope-specific T cells

To characterize the functional T-cell responses elicited against vaccine epitopes, 2 x 10^6^ viable splenocytes in single cell suspension were plated in round bottomed, 96-well culture plates (Corning #3799) in RPMI 1640 medium (Fisher Scientific #72400021) supplemented with heat-inactivated FBS (Fisher Scientific #10500064). Cells were stimulated with 5 μg/mL of synthetic 27mer peptides, peptide pools or were left unstimulated (medium only). Protein transport was inhibited by addition of brefeldin A (GolgiPlug BD #555029) and monensin (GolgiStop BD #554724) two hours after the initiation of stimulation, followed by overnight incubation at 37 °C and 5% CO2. Cells were then washed and incubated with the FcR-blocking anti-murine CD16/32 antibody (Biolegend #101301, 1:50) at 4 °C for 10 minutes, followed by staining for murine CD3e (FITC, Biolegend #100305, 1:800), CD4 (PE-Cy7, BD #552775, 1:800), CD8 (BV786, BD #563332, 1:250), and viability dye (GloCell^TM^ Fixable Viability Dye, Stem Cell Tech #75010, 1:1000) at 4 °C for 30 minutes. Cells were then fixed (eBioscience #00-8222-49) at 4 °C for 20 minutes and permeabilization (eBioscience #00-8333-56) before staining for intracellular IFNγ (BV650, BD #563854, 1:1050) and TNFα (BV421, BD # 566287, 1:1050) at 4 °C for 30 minutes. Stained cells were acquired on FACS Celesta (BD) and frequencies of cytokine-producing CD4+ and CD8+ T cells were determined in FlowJo Software (version 10.8.0). The gating strategy is presented in Figure S4B.

#### B16F10 tumor immune phenotyping

B16F10 immune phenotyping experiments were conducted as described previously^37^. Tumor digest single cell suspensions were incubated with FcR-blocking anti-CD16/32 antibody (Biolegend #101301, 1:50) for 10 min at 4 °C followed by antibody surface staining for CD45.2 (PerCP-Cy5,5 BD #552848, 1:400), CD3ε (FITC, Biolegend #100305, 1:800), CD4 (PE-Cy7, BD #563933, 1:800), CD8 (BV786, BD #563332, 1:250), viability dye (GloCell™ Fixable Viability Dye, Stem Cell Tech #75010, 1:1000) and NK1.1 (BV605, Biolegend #108753, 1:200) for 30 min at 4 °C. Cells then underwent fixation and permeabilization through addition of Foxp3 Fixation/Permeabilization solution prepared by mixing one part of the concentrate with three parts of the diluent (Foxp3/Transcription Factor Staining Buffer Set, eBioscience™ #00-5523-00) for 60 min at 4 °C. Cells were then stained intranuclearly for FoxP3 (PEeFluor610, Thermo Fischer, #61-5773-82, 1:400) in permeabilization buffer (eBioscience™ #00-8333-56) for 30 min at 4 °C. Flow cytometry was performed on FACS Celesta (BD) and the frequencies of the immune populations were determined in FlowJo software (version 10.8.0). The gating strategy is presented in Figure S5.

#### Patient stratification based on EVE burden

RNA-seq of tumor biopsies were retrieved from Liu et al.^32^, van Allen et al.^33^, Riaz et al.^34^ and Mariathasan et al.^35^. Transcript expression levels were quantified as described above (Section: Quantification of EVE expression) and the EVE burden defined as the total number of EVE transcripts with expression levels above 1 transcripts per million (TPM). Study-wise medians were applied as thresholds to group the patients into EVE-High and EVE-Low strata.

WES of matched tumor and healthy tissue biopsy was retrieved from Liu et al.^32^, van Allen et al.^33^ and Mariathasan et al.^35^. The WES reads were mapped to the human reference genome (GRCh38) with BWA-MEM^70^ v0.7.17-r1188. Read duplicates were marked and removed using SAMtools^71^ v1.13 followed by somatic variant calling with GATK-Mutect2^72^ v4.2.2.0. Somatic variants were annotated using the variant effect predictor^73^ v104 and the Ensembl GRCh38.104 annotation reference. Furthermore, annotated somatic variants were retrieved from the supporting material of Riaz et al.^34^. The tumor mutational burden (TMB) was computed as the number of missense somatic variants with a variant allele frequency exceeding 5%. Study-wise medians were applied as thresholds to group the patients into high and low mutational burden strata.

Survival analysis was conducted using the python package lifelines v0.27.3^74^.

#### Pan cancer analysis

Matched TCGA^75^ deposited RNA-seq and Mutect2 somatic variants calls were retrieved. The EVE burden was computed as described above, while the Mutect2 variant calls were subset to the coding regions followed by annotation with the variant effect predictor^73^ v104.

#### Healthy tissue analysis

RNA-seq of healthy tissue samples were retrieved from the GTEx project^76^ and expression levels computed as described above. TPM values were transformed using the Yeo-Johnson approach and projected using the Uniform Manifold Approximation and Projection (UMAP). Clustering was done with agglomerative hierarchical clustering of the UMAP vectors using Euclidean distance and average linkage. Clusters were assessed using the Silhouette score, Adjusted Mutual Information and Purity score. Tissue centroids were defined by the tissue-wise median of the UMAP vectors and tissue distances computed using the same clustering framework. For each tissue except testis, the fraction of samples supporting expression of a given EVE (TPM>1) was computed. If the maximum of these fraction was less than 0.05, the EVE was considered expressed a safe vaccine target. The same procedure was applied for the Ensemble v104 annotated protein-coding genes. The MAGE family (MAGE-A1/A2/A3/A4/A6/A9/A19/A12/C1/C3) was used as a comparator for tumor specificity.

### QUANTIFICATION AND STATISTICAL ANALYSIS

For immunopeptidomics data, incorrect peptide identification was estimated based on the target-decoy search strategy described previously^77^ where the false discovery rate (FDR) variable indicates the proportion of likely incorrect results contained at any given export file. All searches were exported with an FDR of 5%. For immunopeptidomics data, incorrect peptide identification was estimated based on the target-decoy search strategy described previously^70^ where the false discovery rate (FDR) variable indicates the proportion of likely incorrect results contained at any given export file. All searches were exported with an FDR of 5%.

For preclinical data, GraphPad Prism 9 for Mac OS X was used for graphing, statistical analyses, and tools. Data were subjected to Kolmogorov-Smirnov test for normality (alpha = 0.05). Parametric data were analyzed by ordinary ANOVA with Sidak’s multiple comparison correction. Non-parametric data were analyzed by Mann-Whitney test (if two comparisons) or Kruskal-Wallis test with Dunn’s multiple comparison correction (if more than two comparisons). For all preclinical results, the following levels of statistical significance are applied: *p < 0.05, **p < 0.01, ***p < 0.001, ****p < 0.0001.

The lifelines python package v0.27.3^74^ was used for survival analysis and survival statistics (log-rank p-values and hazard ratios). Clustering performance metrics and correlation coefficients were calculated using SciPy^78^ v1.9.3. Average precision scores were calculated using scikit-learn^68^ v1.1.3.

## ACKNOWLEDGEMENTS

We would like to acknowledge Mads Lausen Nielsen, Anne Lund and Marianne Blirup Jensen (Evaxion Biotech) for their assistance with *ex vivo* experiments. Furthermore, we wish to thank Henriette Løwe Press, Heidii Mazur and Rikke Sølberg (Evaxion Biotech) for their excellent execution of preclinical studies and meticulous monitoring of the mice well-being. NJB is a fellow of the Lundbeck Foundation (R272-2017-4040) and acknowledges funding from Aarhus University Research Foundation (AUFF-E-2018-7-14), and the Novo Nordisk Foundation (NNF21OC0071483). AWP is a National Health and Medical Research Council of Australia Investigator Fellow. PF was granted the Victorian Cancer Agency Fellowship.

## AUTHOR CONTRIBUTIONS

CG, MAP, PG, EJL, BR and TT conceived the study and designed the experiments. CG, MAP, PG and TT wrote the manuscript. CG, PG, MS, JA and TT performed computational analysis. MAP, SHR, KP, PF, SC and SF performed experiments. DKK, NJB, JVK and AWP provided further intellectual input. All authors read and approved the manuscript.

## DATA AVAILABILITY

- Raw RNA sequencing data generated in this study have been deposited in the NCBI GEO database under accession number GSE223515.
- Raw mass spectrometry proteomics data have been deposited in ProteomeXchange Consortium via the PRIDE^57^ partner repository under accession number PXD040085 for CT26 and PXD040165 for B16F10 datasets.
- Accessions for datasets used in this study include: SRP011540, SRP094781, SRP012682

## CODE AVAILABILITY

- Code used to generate results and figures included in this manuscript is available on GitLab.com: https://gitlab.com/evaxion/publications/eve-antigen-source

