## Supplementary Figures for "Endogenous viral elements constitute a complementary source of antigens for personalized cancer vaccines"

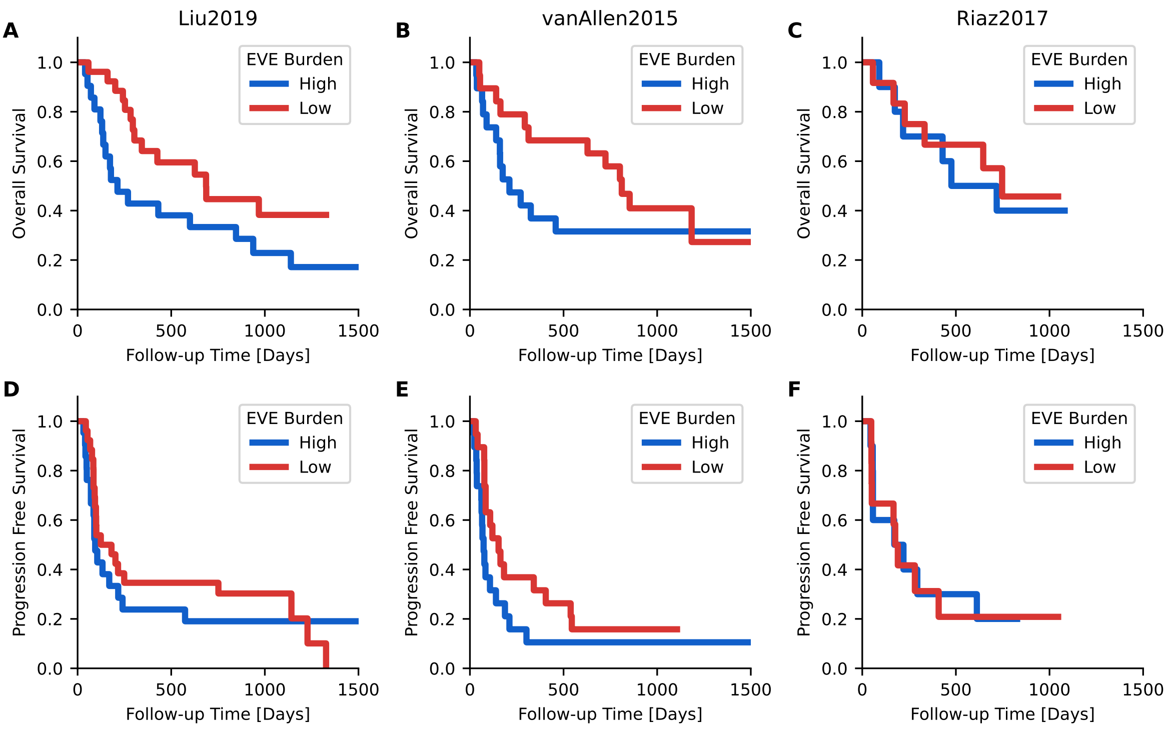


**Figure S1: EVE antigen burden stratification of melnaoma patients receiving checkpoint inhibitors.** (A) Overall survival analysis of EVE strata for Liu et al. (2019) [1]. (B) Overall survival analysis of EVE strata for van Allen al. (2015) [2]. (C) Overall survival analysis of EVE strata for Riaz et al. (2017) [3]. (D) Progression-free survival analysis of EVE strata for Liu et al. (2019) [1]. (E) Progression-free survival analysis of EVE strata for van Allen al. (2015) [2]. (F) Progression-free survival analysis of EVE strata for Riaz et al. (2017) [3].

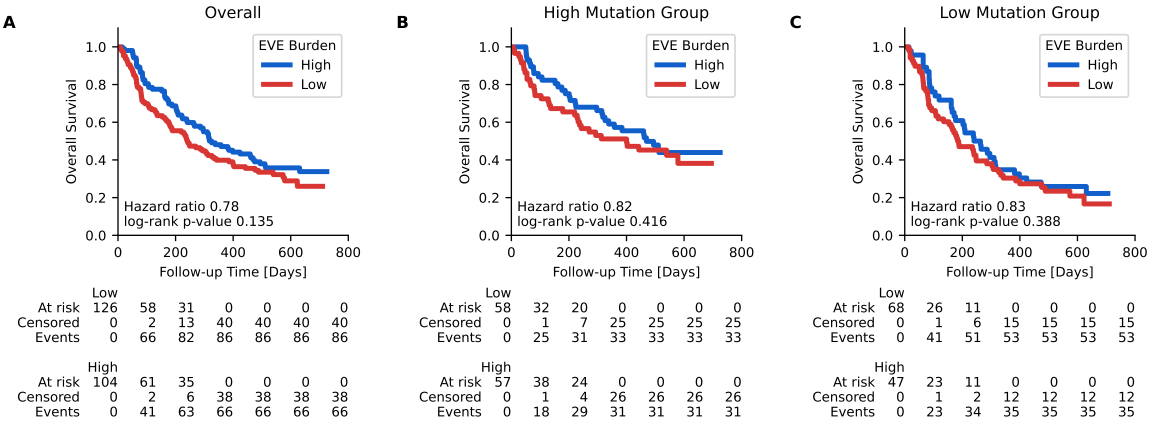


**Figure S2: EVE antigen burden stratification of patients from the Mariathasan et al. (2018) [4] study.** (A) Overall survival analysis of EVE strata for all patients. (B) Overall survival analysis of EVE strata for patients assigned the high mutation group. (C) Overall survival analysis of EVE strata for patients assigned the low mutation group.

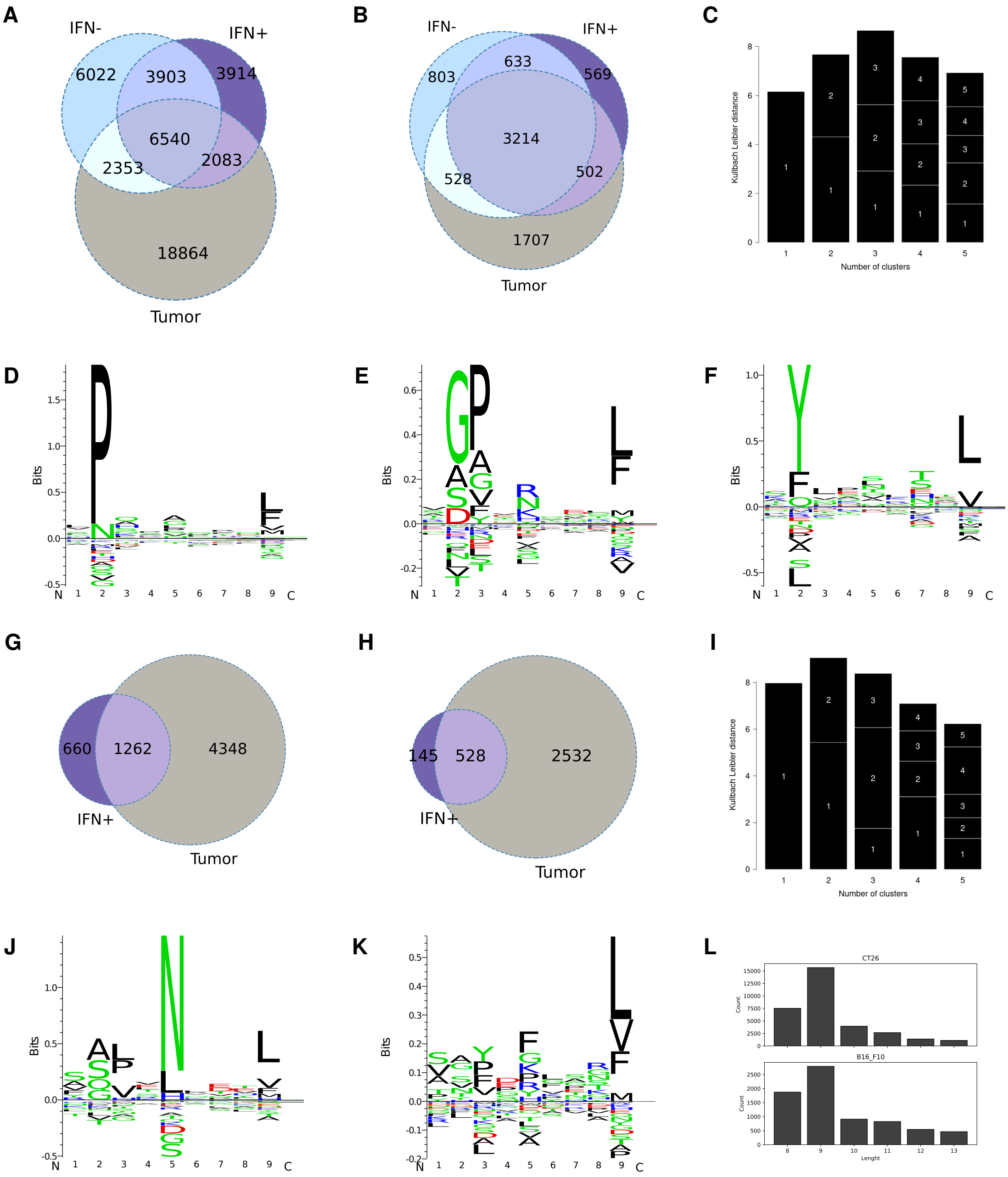


**Figure S3: Overlap and characteristics of murine MHCI ligands identifyed by immunopeptidomics.** (A) Pre filtering overlap of CT26 MHCI ligands for unstimulated cell culture [IFN-], IFNy treated cell culture [IFN+] and *in vivo* grown tumors. (B) Post filtering overlap of CT26 MHCI ligands for unstimulated cell culture [IFN-], IFNy treated cell culture [IFN+] and *in vivo* grown tumors. (C) Motif deconvolution optimization of CT26 immunopeptidomics using GibbsCluster [5]. (D-F) Sequence logos based on the optimal motif deconvolution of CT26 immunopeptidomics, respectively representing H2-Ld, H2-Dd and H2-Kd. (G) Overlap of B16F10 MHCI ligands for IFNy treated cell culture [IFN+] and *in vivo* grown tumors. (H) Overlap of B16F10 MHCI ligands for IFNy treated cell culture [IFN+] and *in vivo* grown tumors. (I) Motif deconvolution optimization of B16F10 immunopeptidomics using GibbsCluster [5]. (J-K) Sequence logos based on the optimal motif deconvolution of B16F10 immunopeptidomics, respectively representing H2-Db and H2-Kb. (L) Length distributions of respectively the CT26 and B16F10 MHCI ligands.

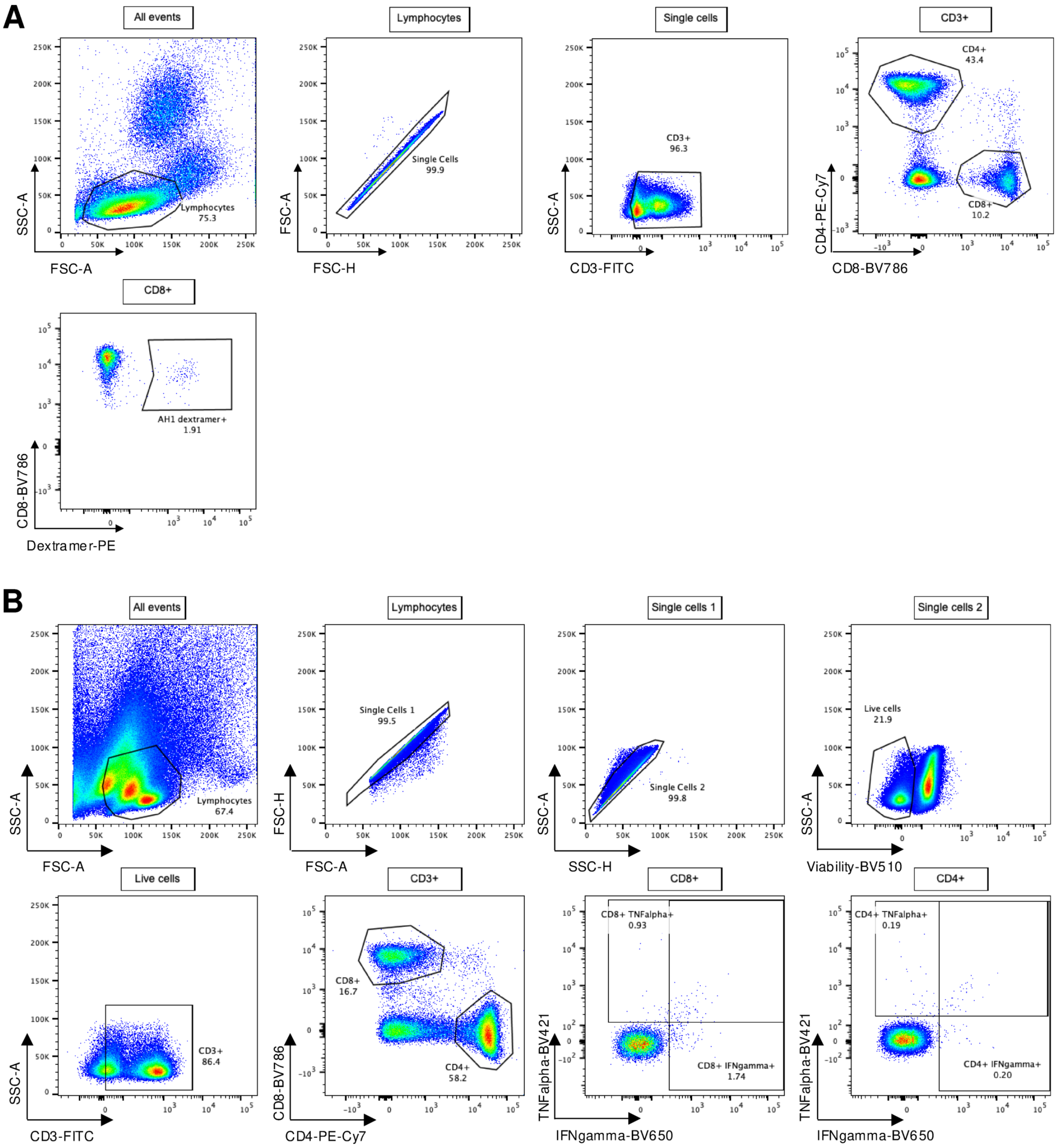


**Figure S4: FACS gating strategies for MHC class I dextramer staining and for peptide re-stimulation and Intracellular Cytokine Staining (ICS).** (A) MHCI dextramers comprised H2-Ld molecules loaded with the restricted minimal peptide SPSYVYHQF (AH1) and antibodies specific for murine CD3, CD4 and CD8 markers were used to stain tail vein blood. Dextramer-positive gate was set based on fluorescence minus one (FMO) control and staining of blood from age-matched naïve mice. (B) Splenocytes were stained with viability dye and antibodies specific for murine CD3, CD4, CD8, IFNy and TNF-alpha markers. Single cytokine-positive gates were set based on non-stimulated samples

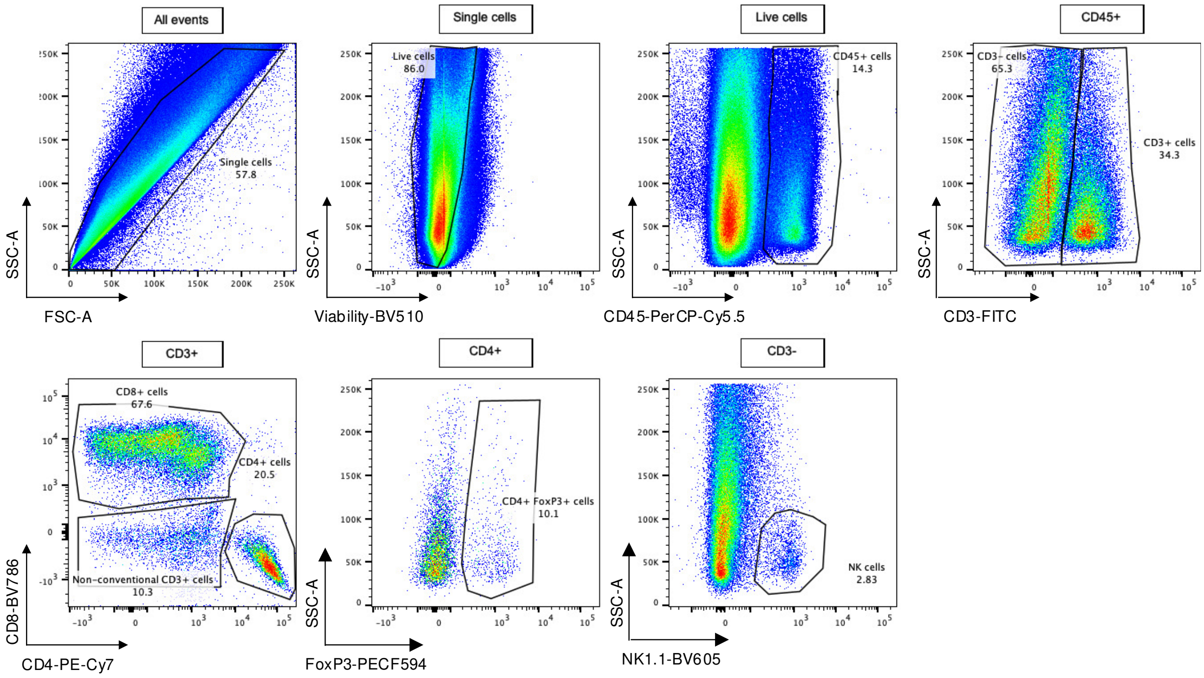


**Figure S5: FACS gating strategy for immune cell phenotyping of digested B16F10 tumors.** Single cell suspensions of B16F10 tumor digests were stained with viability dye as well as antibodies specific to CD45.2, CD3$\epsilon$, CD4, CD8, FoxP3 and NK1.1. Gates were determined based on splenocytes.
